# Comparing Neural Correlates of Memory Encoding and Maintenance for Foveal and Peripheral Stimuli

**DOI:** 10.1101/2023.10.09.561562

**Authors:** G. Kandemir, C.N.L. Olivers

**Author notes:** Address correspondence to: Güven Kandemir, Institute for Brain and Behaviour, Vrije Universiteit Amsterdam Van der Boechorststraat 7, 1081 BT Amsterdam, The Netherlands.

## Abstract

Visual working memory is believed to rely on top-down attentional mechanisms that sustain active sensory representations in early visual cortex. However, both bottom-up sensory input and top-down attentional modulations thereof have been shown to be biased towards the fovea, at the expense of the periphery, and initially peripheral percepts may even be assimilated by foveal processing. This raises the question whether and how visual working memory differs for central and peripheral input. To address this, we conducted a delayed orientation recall task in which an orientation stimulus was presented either at the center of the screen or at 15° eccentricity to the left or right. Response accuracy, electroencephalographic (EEG) activity and gaze position were recorded from 30 participants. Accuracy was slightly but significantly higher for foveal versus peripheral memories. Orientation decoding analyses of the EEG signals revealed a clear dissociation between early sensory and later maintenance signals. While sensory signals were clearly decodable for the central location, they were not for peripheral locations. In contrast, maintenance signals were decodable to an equal extent for both foveal and peripheral memories, suggesting comparable top-down components regardless of eccentricity. Moreover, while memory representations were initially spatially specific and reflected in voltage fluctuations, later in the maintenance period they generalized across locations, as emerged in alpha oscillations, thus revealing a dynamic transformation within memory. Moreover, these transformed representations remained accessible through sensory impulse-driven perturbations, unveiling the underlying memory state. The eccentricity-driven dissociation between disparate sensory and common maintenance representations indicates that storage activity patterns as measured by EEG reflect signals beyond primary visual cortex.

## Introduction

Visual working memory is crucial in that it enables the cognitive system to flexibly and temporarily maintain visual information in the absence of ongoing stimulation. One prominent account, known as the *sensory recruitment hypothesis*, proposes that visual working memory is implemented through top-down activation of sensory representations in visual cortex (Awh & Jonides, 2001; D’Esposito, 2007; D’Esposito & Postle, 2015; Postle, 2006; Serences, 2016; Sreenivasan et al., 2014). However, visual cortex is far from uniform, as a considerably smaller proportion of neural tissue is dedicated to peripheral vision than to central, foveal vision, resulting in concomitant information loss, but also in changes in functionality (Dumoulin & Wandell, 2008; Horton & Hoyt, 1991; Rosenholtz, 2016; Rovamo & Virsu, 1979; Strasburger et al., 2011). Little is known what the impact of this non-uniformity is on visual working memory. To address this, the present study contrasted memory for peripherally and foveally presented stimuli using behavioral and neurophysiological measures.

Differences between central and peripheral vision start in the retina, as distribution and density of receptor types vary with eccentricity (Curcio et al., 1990). Moreover, for peripheral signals there is stronger convergence onto subsequent ganglion cells, resulting in larger receptive fields (Curcio & Allen, 1990). This scheme is continued in the brain, where stronger pooling of signals along the visual hierarchy results in fewer neurons and even larger receptive fields for peripheral vision compared to central vision (Dumoulin & Wandell, 2008; Horton & Hoyt, 1991; Rovamo & Virsu, 1979). As a consequence, increased eccentricity comes with a decline in acuity and contrast detection, as well as increased interference from visual crowding (Levi, 2008; Pelli, 2008; Pelli & Tillman, 2008; Strasburger et al., 2011;Whitney & Levi, 2011). Interestingly, recent studies indicate that visual crowding may behave differently for representations maintained in visual working memory than for sensory representations of the same stimuli when present in the visual array (Harrison & Bays, 2018; Yörük et al., 2020). This would suggest that visual working memory is not simply sustained activation of the same representations as used for visual perception. This may mean that sensory and mnemonic signals tap into different representations after all, either between or within visual layers. Consistent with the latter, feedback signals associated with visual working memories have been shown to tap into different layers of V1 than the input signals (van Kerkoerle et al., 2017).

Importantly, neither may such top-down feedback signals be uniform across the visual field. According to the sensory recruitment and similar hypotheses, memory representations in early visual cortex are being activated through top-down feedback connections. More often than not, these feedback signals have been equated with attentional mechanisms, in the sense that active working memory maintenance is seen as “internal attention” (e.g., Awh & Jonides, 2001; Chun, 2011; D’Esposito & Postle, 2015; Kiyonaga & Egner, 2013; Olivers, 2008; Postle, 2006), and conversely, information maintained in working memory is necessarily information attended (e.g., Desimone & Duncan, 1995; Olivers et al., 2006; Soto et al., 2008). Interestingly, evidence suggests that peripheral vision is not only disadvantaged in terms of the input signal, but also by attention. For instance, it has been shown that peripheral stimuli are more frequently missed under conditions of cognitive load induced by foveal stimuli, resembling a tunnel-vision effect (Ball et al., 1993, Ikeda & Takeuchi, 1975; Williams 1988; 1989; Ringer et al., 2016). Similarly, detection and discrimination performance diminish with eccentricity even when cortical differences are compensated for by cortical magnification (Carrasco et al., 2003; Staugaard et al., 2016), while attentional resolution appears further limited in the periphery above and beyond visual acuity (Intriligator & Cavanagh 2001). Both saliency- and relevance-driven attentional selection mechanisms are affected by eccentricity (Van Heusden et al., 2021; 2023a), and when stimuli are placed at different eccentricities, observers prefer to attend to the more central ones (Heusden et al., 2023b; Wolfe et al., 1998). At the neurophysiological level, it has been shown that presenting a stimulus at fixation can be sufficient to extinguish neural responses to a peripheral stimulus (Richmond et al., 1983; see Desimone & Duncan, 1995 for an overview), and that attending to peripheral stimuli leads to relatively stronger modulations of event-related potentials than attending to central stimuli (Neville & Lawson, 1987). Given that attention has been hypothesized to underpin visual working memory maintenance, this raises the important question whether maintenance of peripheral stimuli differs from that of central stimuli. If attention, or any feedback process in general is weaker for the periphery, so may be maintenance.

If maintaining peripheral memory representations is indeed more challenging than foveal representations, there may be mitigating or compensating mechanisms, by leveraging additional, shared representational resources. These common resources can take different forms, with one possibility being the transformation of a peripheral representations into a *global* representation by recruiting similar feature-specific neurons across the visual field. Such globalization has been suggested on the basis of studies showing that information on memoranda can be reconstructed from non-stimulated portions of early visual cortex (Ester et al., 2009; Serences et al., 2009). Second, peripheral representations may be *fovealized*, such that specifically neural sensory tissue that is normally dedicated to the fovea is being recruited to aid in, or even replace the representation of peripherally presented memoranda. Foveal patterns of fMRI activity have been demonstrated to carry information on peripheral stimuli (Williams et al., 2008), and foveal disruption has been shown to hinder performance in peripheral discrimination tasks (Chambers et al., 2013; Fan et al., 2016; see also Stewart et al., 2020). A third possibility is that the mnemonic representation – whether originally peripheral or foveal – is being transformed into a common representation that is less sensory, and more *functional* in nature. Such functionalization would then serve the prospective purpose of the memory, which, depending on the test, could for example be a more spatial, attentional representation, or an intentional, motor representation (Cochrane & Green, 2023; Mostert et al., 2018; Myers et al., 2017; Olivers & Roelfsema, 2020; van Ede & Nobre, 2023). Importantly, while these three options (*globalization*, *fovealization*, and *functionalization*) differ, they all predict a *generalization* between central and peripheral representations, as well as between different peripheral representations.

To investigate if, and how, central and peripheral stimuli may be differentially represented in visual working memory, and how this may dynamically change over time, we conducted an EEG experiment in which we asked human observers to memorize on each trial the orientation of a single Gabor stimulus. Figure 1 illustrates the task and procedure. Crucially, the stimulus was presented either at the center of vision or in the periphery at 15° eccentricity to the left or right of fixation. In order to increase the chances of being able to trace the memory signal also during activity-silent periods, we also included two memory “pinging” or “perturbation” episodes, which involved an array of white disks that were presented during maintenance phase, serving as impulse signals and which have previously been shown to boost memory decoding when activity patterns have decayed (Kandemir & Akyürek, 2023; Stokes, 2015; Wolff et al., 2017; 2020). We were specifically interested in whether and to what extent memory encoding and maintenance signals associated with peripheral items were weaker than for central items. To this end, we sought to decode orientation-specific patterns of activity in both raw voltage and frequency-decomposed signals within alpha band, for central and peripheral positions. If top-down maintenance signals indeed weaken with eccentricity, we should observe a stronger decline in the strength of location-specific orientation classification for peripheral stimuli. To foreshadow, this is not what we found. Second, we were interested if and to what extent shared representations were employed during memory maintenance. Whether due to globalization, fovealization, or functionalization, shared representations predict generalization. To test this prediction, we trained and tested our classifiers across the different stimulus locations, and assessed whether, when, and for what type of signal the classification generalized between central and peripheral memoranda. Again to foreshadow, we did observe such generalization, but only relatively late into the delay period, and only in the alpha band – consistent with an attentional or intentional preparation for the test.

**Figure 1.**
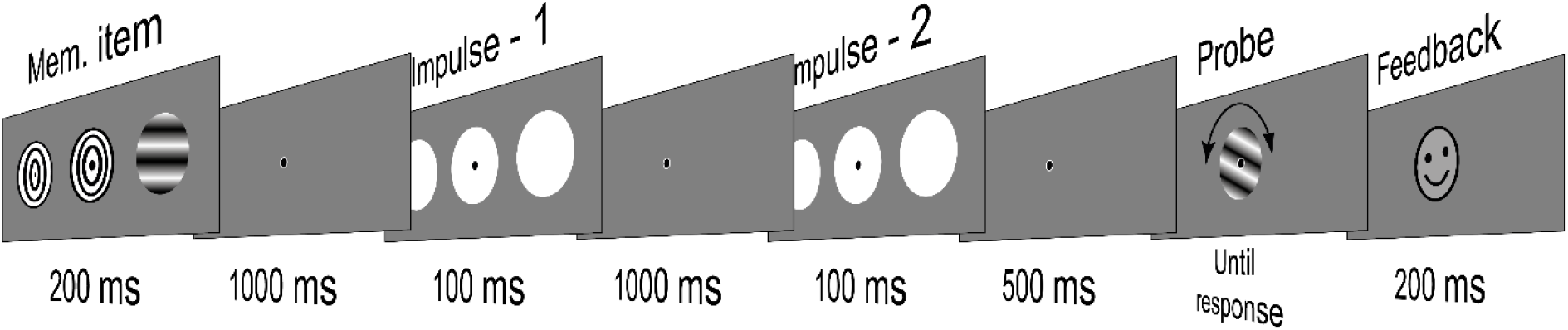
An outlook of a single trial design. Note. In each trial, one Gabor patch with varying orientations was presented at one of three possible locations, whereas other locations were filled with concentric circles forming a bulls-eye symbol. At the end of the trial, orientation was reported by rotating the random probe via PC compatible controller.

## Method

### Participants

Thirty healthy young adults volunteered to participate in the study (23 female, M_Age_ = 20.0, s.d._Age_ = 1.81) in return for 13 euros/hr. The sample size was estimated from earlier studies using the impulse-based memory perturbation technique (e.g., Wolff et al., 2017). Participants were informed about the general aim of the study and the procedure both verbally and in text prior to participation. Participants provided written consent to declare their willingness to participate, their compliance with the instructions, and their agreement with the data storage and data sharing policies. All participants except one completed entire session with 2880 trials, whereas for one participant 96 trials were lost due to a technical error The study followed the general protocol approved by the Scientific and Ethics Review Board of the Faculty of Behavioral and Movement Sciences, VU University Amsterdam (protocol number VCWE-2021-173).

### Stimuli and Apparatus

A gray background (RGB = 128, 128, 128;, 76.3 cd/m^2^) was maintained throughout the experiment. A fixation dot consisting of a black dot (0.3° diameter) surrounding a white circle (0.25° diameter) was displayed at the center of the screen throughout a trial. An oriented Gabor stimulus served as the memory item on each trial (81.2 cd/m^2^). The orientation was randomly selected from a set of 24 discrete equidistant angles ranging from 3.75° to 176.25° (7.5° angular distance). The contrast and the spatial frequency of the Gabor stimulus were kept at 1, whereas the phase varied randomly within the range of 1 to 180 for each trial. There were three stimulus locations, one centered on the center point of the screen, and the other two were respectively positioned 15° to the left and right. On each trial, the memory item was displayed on one of these three locations whereas other the two locations were filled by placeholders. The placeholders were concentric black and white circles forming a bullseye. The size of the stimuli, whether memory item or placeholder, was 5° in diameter.

The impulse stimulus corresponded to three white discs (RGB = 255, 255, 255; 306.4 cd/m^2^) centering the same locations as the memory item and the placeholders. Each disc had a diameter of 7.5°. In the response screen, a manipulable single Gabor stimulus with a diameter of 5° was displayed at the center of the screen. The initial orientation of the displayed Gabor was randomly drawn on each trial. The contrast and the spatial frequency of the stimulus were set to 1. The phase varied between 1 to 180 and was randomly selected on each trial. The participant then rotated the orientation with the help of a joystick to match their memory. A feedback consisting of a smiley was displayed to encourage participants for higher performance. The smiley size was 5° in diameter. Responses with an absolute deviation of 20° or less relative to the presented memorandum received a happy smiley as a feedback whereas larger deviations received a sad smiley.

All stimuli were created and presented with the Psychtoolbox extension for Matlab (Brainard, 1997; Kleiner et al., 2007). The experiment was displayed on a 23.8 inch (60,452 cm on diagonal) ASUS ROG Strix XG248Q monitor with a resolution 1920 by 1080 pixels and a refresh rate of 240 Hz. The responses were collected by a windows compatible Xbox controller with a USB cable, where the left joystick was used to rotate the probe and A button was used to submit the response. Throughout the experiment, simultaneous eye position and pupil size of the left eye was recorded at 1000 Hz sampling rate by desktop mounted Eyelink 1000 Plus.

### Procedure

The experiment consisted of 2880 trials distributed equally over five consecutive sessions, each lasting approximately 65 minutes. During a session participants were seated in a dimly lit room at 60 cm distance from the monitor with their head placed on the chinrest. Each session was divided into 24 blocks consisting of 24 trials. Between sessions, participants took breaks with self-regulated durations to combat fatigue and within a session participants could also rest between blocks during which their performance in that session so-far was displayed. Within a block, trials were automated such that following the submittance of a response, next trial began. A single trial is illustrated in Figure 1.

Each block began by a SPACE bar press. A ‘Get Ready’ message was then displayed in Arial font in size 12 for 400 ms and a blank screen was displayed for 600 ms. This was followed by the presentation of the fixation dot, which stayed on the screen for the remainder of the trial. After 700 ms relative to fixation dot onset, the memory item and the placeholders were presented for 200 ms. A delay period of 1000 ms followed with only the fixation dot on display. Next, the first impulse stimulus was presented for 100 ms duration and another delay followed for 1000 ms. The second impulse was presented for 100 ms duration, followed by another 500 ms delay. Next, the response screen was displayed, during which the orientation of the probe could be manipulated by the participants by tilting the joystick on the controller in order to report the orientation of the memory item. Pressing button A on the joystick submitted the response. Following the response, a blank screen was displayed for 200 ms which was followed by the feedback, which was presented for 200 ms. The fixation cross for the next trial then followed after a delay with a random jitter of 200 – 500 ms.

Participants were instructed to maintain their gaze fixated on the central dot throughout the entire trial until the response screen. In case of too frequent deviation, participants were given renewed verbal instructions at the end of the block. Eye movements were recorded simultaneously with the EEG. The eye-tracker was re-calibrated using a 5-dot calibration method before the beginning of a block each time the head was moved off the chinrest. Participants were encouraged to respond as fast and accurately as possible and blink only during the response stage.

### EEG Acquisition and Preprocessing

The EEG was recorded from 64 channels deployed according to the international 10-20 system via BioSemi Active 2 system. Four external electrodes placed below and above right eye (Vertical) and lateral to external canthi (Horizontal) recorded bipolar EOG. Two additional electrodes placed on the mastoids would serve as offline reference.

Offline, the data was first re-referenced to the average of the recordings from the two electrodes on the mastoids. Next, the data was downsampled to 500 Hz. By using the freely available Matlab extension EEGLab (Delorme & Makeig, 2004), a bandpass filter (0.1 Hz high – 40 Hz low) was applied. The EEG trial data was then divided into epochs relative to displayed stimuli (-200 ms to 1200 ms relative to Memory item; -200 ms to 1100 ms relative to Impulse 1; -200 ms to 600 ms relative to Impulse 2 and – 200 ms to 300 ms relative to Probe). The summary statistics corresponding to the variance and maximum values on each channel in each trial were first visualized by using the freely available Matlab extension Fieldtrip (Oostenveld et al., 2011). Noisy electrodes were replaced by spherical spline interpolation. On average 1.5 electrodes were removed per participant, where in turn 23 % of these were part of the 19 posterior electrodes used in the main decoding analyses.

Summary statistics were computed and then trials with a variance above 2000 were first marked as candidates of removal due to eye movements. Next, entire sets of epochs were visualized with EEGlab and inspected for muscle artefacts and blinks. Epochs with artefacts were manually marked and excluded from further EEG analyses with the exception of the Probe epoch. In this latter epoch, no trials were excluded due to eye-movement related artefacts like blinks and/or saccades because all participants were instructed to blink when they were responding. The average removal for epochs were as reported: Memory item epoch ∼14 %, Impulse 1 ∼10 %, Impulse 2 ∼7 % and Probe 2 %.

For the alpha power analysis, non-epoched continuous data were first band-pass filtered (8 Hz high – 12 Hz low). Next, the output at each channel was Hilbert transformed. The absolute values produced by this transformation were epoched the same way as the voltage data. The same electrodes and trials were excluded as in the voltage analyses.

### Behavioural Analyses

All existing trials were included in the behavioural analyses. The memory quality was assessed by standard mixture modelling, using the freely available Mem_Toolbox (Suchow et al., 2013). For each participant, stimulus and error data were entered in the model as angular values. Two parameters were estimated, namely the guess rates (corresponding to the estimated proportion of responses belonging to the underlying univariate distribution), and the standard deviation of the gaussian fit to the response error distribution. The estimated standard deviation reflects the width of the error distribution relative to the actual stimulus, and therefore it is a measure of precision, such that higher estimates address worse memory quality. Next, estimated guess rates and standard deviation of participants were analysed as a group with Repeated measures ANOVA, using the freely available statistics program JASP (JASP Team, 2023), where informative, pairwise comparisons were also calculated (*p* < 0.05).

### Multivariate Analyses

Time course orientation decoding was conducted using data from 19 posterior electrodes (PO7, PO3, POz, Pz, P1, P3, P5, P7, P9, Oz, P2, P4, P6, P8, O1, P10, PO8, PO4, O2) for voltage data and alpha power. The decoding was conducted separately for orientations presented at the same location unless stated otherwise and for peripheral items, decoding on Left and Right were averaged at each time point. First, the trial-specific orientations were grouped into 8 equidistant bins, each covering a range of 22.5°. As a part of a cross-validated decoding procedure, the data was partitioned into 8 folds with stratified sampling, and the data of 7 folds were distributed to respective bins as the training set. The covariance matrix was computed by a shrinkage estimator (Ledoit & Wolf, 2004) using the entire data in the training set. For the training set, the trial-count in each bin was equalized via randomly sampling these trials to the trial count of the smallest bin. The EEG data of trials in each bin was averaged to form a representation of the orientation reflected by each bin. The averaged data was then convolved with a half cosine basis set raised to the 15^th^ power in order to reduce noise and pool information across similar orientations (Myers et al., 2015). We then took the Mahalanobis distance as the quantification of similarity (De Maesschalck et al., 2000), as estimated between each trial in the left-out fold and the averages of 8 bins. These 8 distance measures estimated for each trial were mean-centered and the sign was reversed so that higher values would indicate higher similarity. Re-ordering of distance measures to center the target item each time and then averaging these values provided similarity matrices. Convolution of the distance values at each time point with a cosine function yielded the decoding accuracy (in arbitrary units). This procedure was applied to all time points within an epoch. The procedure was repeated 8 times so that all trials took part in the test set. The entire procedure was repeated over three orientation spaces so that each orientation in a bin was closest to the bin center once (0° – 157.5°; 7.5° - 165° and 15° - 172.5°). In order to avoid sampling bias, the decoding over each orientation space was also repeated 100 times, resulting a total of 300 repetitions. The decoding accuracy measures and the distance values were averaged over all repetitions and trials for each participant. The time course data was smoothened by a gaussian filter (kernel = 2, 16 ms). The statistical analyses were conducted over group data.

In case of non-circular stimuli here specifically the decoding of presentation locations, the analysis was identical to the analysis of circular data up to the calculation of the Mahalanobis distances with 8-fold cross-validation procedure. The measure of decoding accuracy was then the mean distance difference between the target item and all other possible options, indicating the size of discriminability in that trial at that time point. These distance differences were treated the same way as the cosine-convoluted Mahalanobis distances, by means of averaging over repetitions, trials and the means were smoothened by a gaussian filter (kernel = 2, 16 ms). Statistical analyses were also identical to circular stimuli.

Temporal generalization matrices were formed by a similar 8-fold cross-validation procedure as the time course decoding. The main distinction was the usage of condition-specific patterns formed at each time point to test all possible time points in that epoch. If a generalization of the neural code across two independent conditions was assessed (e.g., distinct spatial locations) the entire data in one condition was used as the training set, whereas the data from the other condition served as the test set. The output matrices reflected time-specific training and testing at the diagonal whereas off-diagonal accurate decoding indicated temporal generalization of the neural code at that specific time to a larger time window.

In order to assess the decoding results over a certain time period, trial-specific memory content was decoded using de-meaned residuals of the raw EEG data within a specified time window. For this, first data in each trial on each channel within the designated time window was averaged and subtracted in order to serve as a baseline. Next, residual data was downsampled to 100 Hz by calculating a moving mean over 10 ms windows, yielding 30 temporal features. These features were pooled over electrode space, forming 570 spatio-temporal features for each trial. These spatio-temporal features were used in the 8-fold cross-validated decoding procedure just like time course analyses described earlier, yielding a single decoding accuracy measure for each trial.

A searchlight technique (van Ede et al, 2018) was used to roughly estimate the distribution of the memory-related signal over the scalp by applying multivariate pattern analysis to data from each electrode and two of its closest neighbours. This analysis was similar to a critical time window analysis as the residual data within a 300 ms window in each trial was calculated by subtracting the mean activity. Next, the residual activity was downsampled to 100 Hz by using a moving window with a 10 ms range. Next, an 8-fold cross validated decoding procedure explained earlier was applied to each electrode by using the data on that electrode and two of its closest neighbours. Computed Mahalanobis distances were convoluted by a cosine function, yielding a trial specific decoding accuracy for each electrode. This procedure was repeated over 64 electrodes to cover the entire scalp, and repeated 3 times to cover the entire angular space. The entire procedure was repeated 100 times to account for probable sampling biases. The final output for each participant corresponded to the mean decoding accuracy at each electrode averaged over all repetitions and trials. Visualization was done by projecting the decoding accuracy of significantly contributing electrodes with the Fieldtrip plotting functions (Oostenveld et al., 2011).

### Statistical analysis

Non-parametric sign permutation tests were used to assess the significance of decoding analyses. For each analysis, a null distribution was formed by randomly flipping the sign of the mean decoding accuracy of each participant 100 000 times with a probability of 0.5 and then averaging the output of this permutation. Next, the mean decoding accuracy of the group sample is contrasted to this null distribution and the proportion it belonged to is reported as the p value, where the cut-off was predetermined as *p* < 0.05. For time course analysis, the sign permutation test is applied to each time point with a cluster correction where the cut-off was set to *p* < 0.05. For searchlight analysis, each electrode output is tested individually with a cut-off, *p* < 0.05.

## Results

### Behavioural Results

Overall, participants were quite successful in their recall for memorized orientations in all conditions as indicated by the raw error distributions (See Fig. 2A). The standard mixture model was applied to estimate the guess rate and precision. As expected given the low memory load, the estimated guess rates were low (See Fig. 2B). The distribution of guess rates deviated significantly from normality (*W* = 0.825*, p* < 0.001), so we chose a non-parametric Wilcoxon signed-rank test. The pairwise comparison revealed that the estimated guess rates were lower for Central orientations, relative to Peripheral (*x̅ _Central_* = 1.90 %, *x̅ _Peripheral_* = 4.80 %, *Z _Peripheral vs. Central_* = 3.610*, p* < 0.001, *r_rb_* = 0.755).

**Figure 2.**
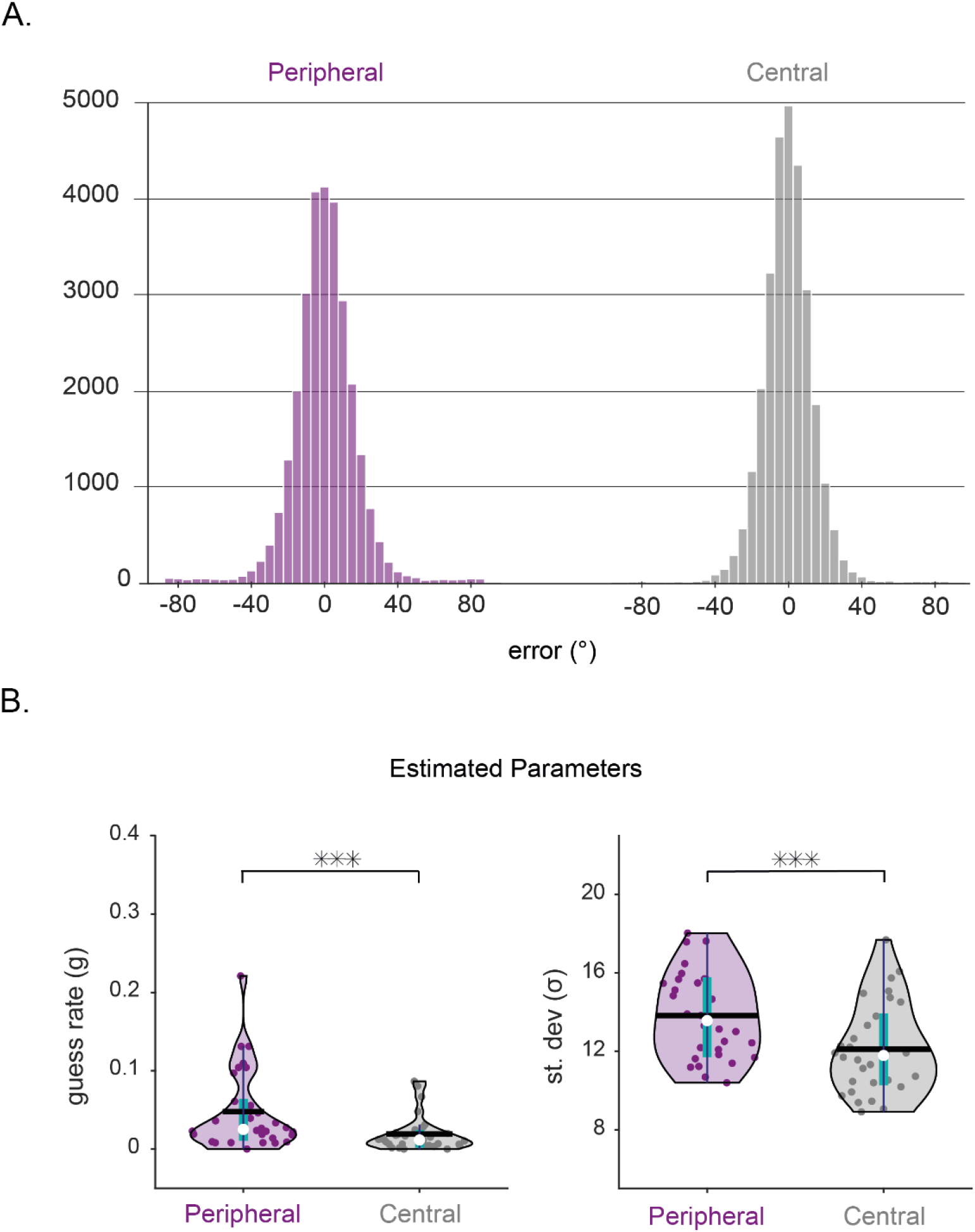
Raw error (A), estimated guess rates (B, left), and estimated standard deviation (B, right) of the mixture model fit for Peripheral (purple) and Central (gray) orientations. *Note.* A. Histogram showing raw errors in the entire data for Peripheral (purple) and Central (gray) orientations. B. Violin plots presenting estimated guess rates (left) and standard deviation (right) as the indication of memory precision, as provided by fitting the standard mixture model. Black solid line marks the mean, whereas the white dot marks the median. The inter quartile range (IQR) is depicted by the green box and the dark blue bar marks 1.5 times the IQR. Asterisks show statistically significant differences between pairs (*p* < 0.001, ***).

For the standard deviation of error distributions the normality assumption held (all *W* < 0.964, *p* > 0.395). A pairwise t-test revealed that when the item was presented at the center, reported memories were more precise relative to when the orientations were presented at periphery (*x̅ _Central_* = 12.11 %, *x̅ _Peripheral_* = 13.82 %, *t _Peripheral vs. Central_ (29)= 7.278, p* < 0.001, *d* = 1.33).

### Multivariate Analysis of Neural Data

#### Decoding Memory Items From Voltage Data

First, to compare the classification strength of Central and Peripheral memory items, we conducted time course analyses on the raw voltage data separately for each location and averaged the decoding accuracy over Peripheral items (See Fig. 3A). The orientations at the center location became successfully decodable shortly after the onset of the visual display and remained traceable until shortly before the end of the epoch (Central, gray, 56 ms – 1080 ms relative to memory item onset, *p* < 0.001, *two-tailed*, *cluster-corrected*). Interestingly, Peripheral orientations were not traceable during the visual presentation. These only became evident after stimulus offset, where decodability gradually became comparable to the central item in terms of both strength and remaining duration (Peripheral, purple, 224 ms – 912 ms relative to memory item onset, *p* < 0.001, *two-tailed*, *cluster-corrected*). Individually, peripheral items yielded weaker but similar results (See Supplements-1, Fig. S1A). Directly comparing classification performance between the Central and Peripheral items yielded stronger decoding for the central item, but this difference was restricted to the early period (Orange area in Fig. 3A, 64 ms to 456 ms relative to memory item onset, *p* < 0.001, *two-tailed*, *cluster-corrected).* Notably, given that representations for peripheral items emerged relatively late and independent of a strong initial sensory response, we argue that these signals do not simply reflect some remaining sensory activity, but are related to working memory maintenance. Furthermore, the lack of any difference in classification strength between peripheral and central items following the stimulation period suggests that differences in behavioural performance reflect differences in encoding phase including sensory effects, instead of differences in memory maintenance. We also decoded eye gaze position as recorded by the eye-tracker to assess whether the current findings can be explained by eye movements (cf. Mostert et al., 2018). While eye movement data allowed for significant above-chance decoding (See Fig. S2 in Supplements-2), this was only the case later into the delay period, and did not overlap with the EEG voltage decoding that we focus on here. We therefore reject the possibility that eye movements induced these effects. We will return to eye movements when discussing the alpha-band decoding.

**Figure 3.**
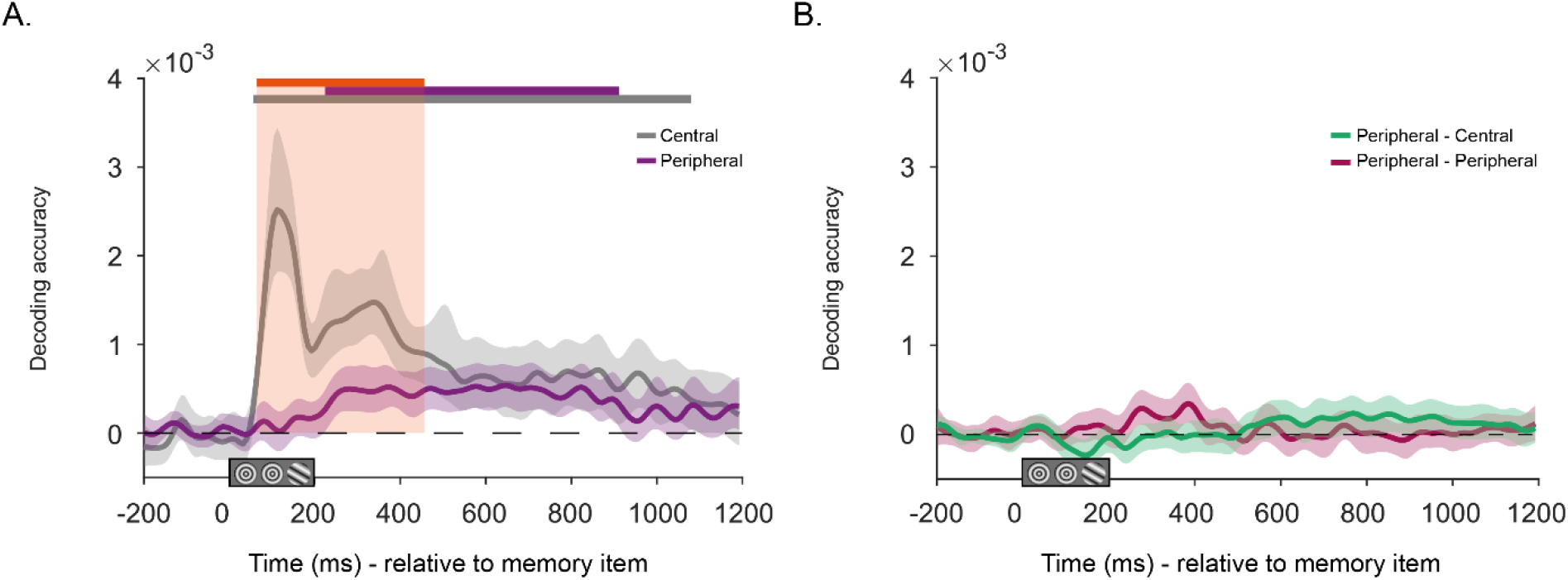
Time-course decoding of raw voltages for trial-specific Peripheral and Central orientations relative to memory item onset (A). Time course generalization of neural code for orientations across locations as reflected in voltage values (B). *Note.* A. Mean decoding accuracy for trial-specific orientations presented at the Central (gray) and Peripheral (purple), as a function of time relative to memory item onset, predicted using the voltage values. The orange rectangle marks the time period with statistically significant difference in decoding accuracy between the central item and peripheral items. B. Mean decoding accuracy for orientations over time relative to first impulse onset, when training and testing on different stimulus locations; reflecting generalization of the neural code across peripheral items (Peripheral - Peripheral, in burgundy), and generalization between peripheral and central items (Peripheral - Central, green), reflected by voltages. A-B. Solid lines show mean decoding accuracy averaged over all trials and participants. The shaded zones around the solid lines mark the 95 % C.I.. Horizontal bars at the top mark statistically significant time periods (*p* < 0.05, *two-tailed, cluster corrected*).

Next, we compared the neural code for orientation representations across the different presentation locations. First, we contrasted the neural code across peripheral locations by training on items presented on one side and testing on items at the other peripheral location (and vice versa). Although Peripheral – Peripheral generalization showed a small increase around 250 – 400 ms period relative to memory item onset, no significant cluster was detected within this epoch (Fig. 3B, Peripheral – Peripheral, burgundy, *p* > 0.05, *two-tailed*, *cluster-corrected*). We also trained on centrally presented items while testing on peripheral items, and vice versa. Neither did this Peripheral – Central generalization analysis yield any significant effects (Fig. 3B, green, *p* > 0.05, *two-tailed*, *cluster corrected*). Note that potential differences in temporal dynamics between different eccentricities could have prevented us from observing any generalizations, when the assessment was restricted to the same time points. Therefore, temporal dynamics were also assessed by including a temporal generalization analysis in which the classifier was trained on items at one peripheral location and tested on another location for all combinations of time points. This again yielded no significant clusters, indicating no generalization for Peripheral – Peripheral (See Fig. S4A in Supplements-3), or for Peripheral – Central decoding schemes (Fig. S4B). These results are consistent with the idea that orientation representations, as traced through raw voltages, remain spatially specific during maintenance.

#### Decoding Memory Items From α Power

Next, we focused on alpha power as a potential alternative carrier of generalized memory information. Fig. 4A shows the decoding results. During sensory encoding, only central orientations were briefly traceable, but this signal soon perished (Fig. 4A, Center, gray, 96 ms to 208 ms, *p* = 0.043). This signal is likely driven by early visual event-related potentials, which tend to fluctuate around the alpha frequency. Orientation information then returned later in the epoch (Fig. 4A, Center, gray, 624 to 1192 ms relative to item onset, *p* < 0.001, *two-tailed*, *cluster corrected*), where it also emerged for Peripheral items (Fig. 4A, Peripheral, purple, 680 ms to 1200 ms relative to memory item onset, *p* < 0.001, *two-tailed*, *cluster corrected*). Crucially, the decoding accuracy in this late period did not differ across central and peripheral items, indicating equivalent top-down maintenance signal strengths independent of eccentricity. Note that here eye movement decoding yielded significant above-chance classification for individual items within the same time period (See Fig. S2 in Supplements-2). To assess whether this explained the alpha decoding (cf. Liu et al., 2022), we correlated the decoding accuracies of EEG alpha power and gaze position at each location and time point, across trials and tested Fisher-transformed correlations against zero. In short, if the alpha decoding is caused by eye gaze, then trials with strong eye decoding should also show strong alpha decoding. However, this analysis yielded no significant clusters (Fig. S3A). To increase power, we also calculated the mean decoding accuracy for both measures within the critical time period (600 – 1200 ms relative to memory item onset), and then tested for the same correlation. This again yielded no significant results, indicating that significant decoding of alpha power was unlikely to be driven by gaze position. Note that this does not mean that the alpha signal and eye gaze cannot reflect the same underlying process – a point that we will return to in the Discussion.

**Figure 4.**
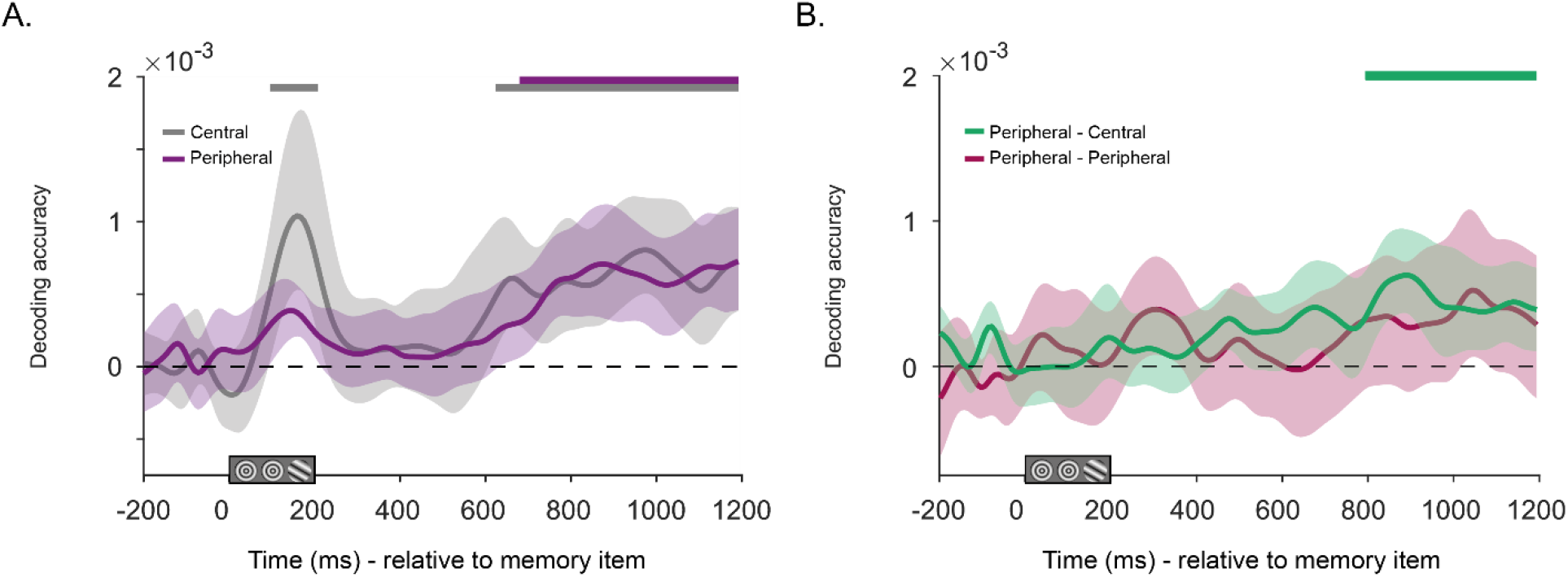
Time-course decoding of absolute change in alpha power for trial-specific Central and Peripheral orientations relative to memory item onset (A). Time course generalization of neural code for orientations across locations reflected by alpha power (B). *Note.* A. Mean decoding accuracy for trial-specific Central (gray) and Peripheral (purple) orientations, as a function of time relative to memory item onset, predicted using the absolute change in the alpha power. B. Mean decoding accuracy for orientations over time relative to memory item onset, when training and testing on different stimulus locations; reflecting generalization of the neural code across peripheral items (Peripheral - Peripheral, in burgundy), and generalization between peripheral and central items (Peripheral - Central, green), reflected by absolute change in alpha power. (A-B) Solid lines show mean decoding accuracy averaged over all trials and participants. The shaded zones around the solid lines mark the 95 % C.I.. Horizontal bars at the top mark statistically significant time periods (*p* < 0.05, *two-tailed, cluster corrected*).

We then progressed to assess the generalization of the alpha power-based signal. To this end, we applied the same train-test approach across items at different locations as we did for the voltage analyses. Importantly, and in contrast to the raw voltage analysis, here training on peripheral items and testing on central items (and vice versa) yielded statistically significant generalization (Fig. 4B, Peripheral - Central, green, 794 ms to 1194 ms relative to item onset, *p* = 0.002, *two-tailed*, *cluster corrected*). Although a similar elevation in the mean decoding accuracy was present for the generalization across the two peripheral locations, this was not statistically significant (Fig. 4B, Peripheral-Peripheral, burgundy, *p* > 0.05, *two-tailed*, *cluster corrected*). These results provide a first indication of a generalized code across central and peripheral memories.

#### Impulse-Driven Memory Decoding

During the memory delay, two impulses were presented sequentially to increase our chance of tracing memory items during activity-silent maintenance periods. First, impulse-driven EEG voltage was decoded for trial-specific orientations, which yielded inconsistent results (See Fig. 1C in Supplements-1), as we observed significant decoding only for the peripheral item on the right side (Right, blue, 216 to 400 ms relative to first impulse onset, *p = 0.019*; Left, red, *p* > 0.05; Center, gray, *p* > 0.05, *two-tailed*, *cluster corrected*). Decoding of orientations following the second impulse also did not yield any significant effects in the voltage data.

In contrast, impulse-related decoding in the alpha band did yield reliable effects (See Fig. 5A). Decoding accuracy increased statistically post-impulse for Central orientations (Central, gray, 112 ms – 1096 ms relative to memory item onset, *p* < 0.001, *two-tailed*, *cluster-corrected*) as well as Peripheral items (Fig. 5A, Peripheral, purple, 48 ms – 1100 ms relative to first impulse onset, *p = 0.001*, *two-tailed*, *cluster-corrected*). Although time precision is reduced by frequency decompositions, the peak accuracy within the epoch was observed within the first 400 ms time period relative to impulse onset, which is consistent with it being caused by the impulse. Crucially, the strength of information did not differ between periphery and center. The decoded signal did not rely on eye movements or gaze position (See Fig. S2B). Given that alpha power was baselined to pre-impulse period, the impulse presentation further boosted the memory-related signal that was already present within the alpha band. Moreover, as in the pre-impulse period, the neural code for orientations generalized across locations also after the impulse. Assessment of both Peripheral - Peripheral and Peripheral – Central generalization yielded significant results (Fig. 5B, Peripheral - Peripheral, burgundy, 112 ms – 256 ms relative to first impulse onset, *p* = 0.046; Peripheral – Central, green, 64 ms – 336 ms, *p* = 0.007; *two-tailed, cluster-corrected*). These results further support the idea further into the delay period, a generalized code emerges in the alpha band. Analyses on the second impulse epoch yielded no additional significant results.

**Figure 5.**
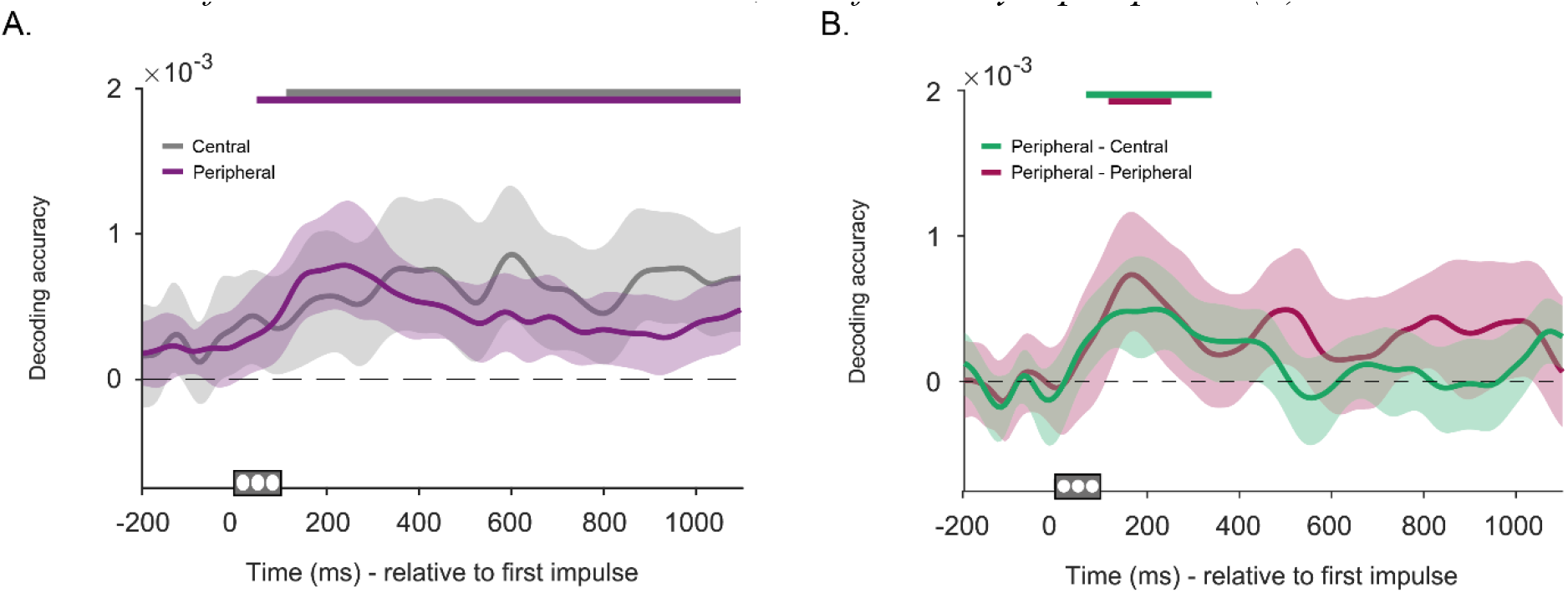
Time-course decoding of absolute change in alpha power for trial-specific Central and Peripheral orientations relative to first impulse onset (A). Time course generalization of neural code for orientations across locations, as reflected by alpha power (B). *Note.* A. Mean decoding accuracy for trial specific orientations presented at the Center (gray) and Periphery (purple) from the absolute change in alpha power as a function of time relative to the presentation of first impulse signal. B. Mean decoding accuracy for orientations over time relative to first impulse onset, when training and testing on different stimulus locations; reflecting generalization of the neural code across peripheral items (Peripheral - Peripheral, in burgundy), and generalization between peripheral and central items (Peripheral - Central, green), as reflected in absolute change in alpha power. A-B. Solid colored lines reflect the mean decoding accuracy. The shaded zones mark the 95 % C.I.. Horizontal bars at the top mark statistically significant time periods (*p* < 0.05, *two-tailed, cluster corrected*).

#### Further Investigations of Memory Encoding for Peripheral and Central Items

As we were quite surprised by the lack of any reliable sensory encoding signals for peripheral items, we conducted a number of post-hoc tests to investigate this further. To assess if there was no orientation information whatsoever for peripheral items in the initial sensory signal, we first sought to increase power by pooling data over the entire stimulus presentation period (0-200 ms relative to stimulus onset; Fig. 6A). This time we could successfully decode peripheral orientations, although the effect remained very weak compared to central items (Left, red, *p* = 0.045; Right, blue, *p* = 0.035; Center, gray, *p* < 0.001; Central-Peripheral, *p* < 0.001; *two-tailed*, *n _perm_* = 100 000).

**Figure 6.**
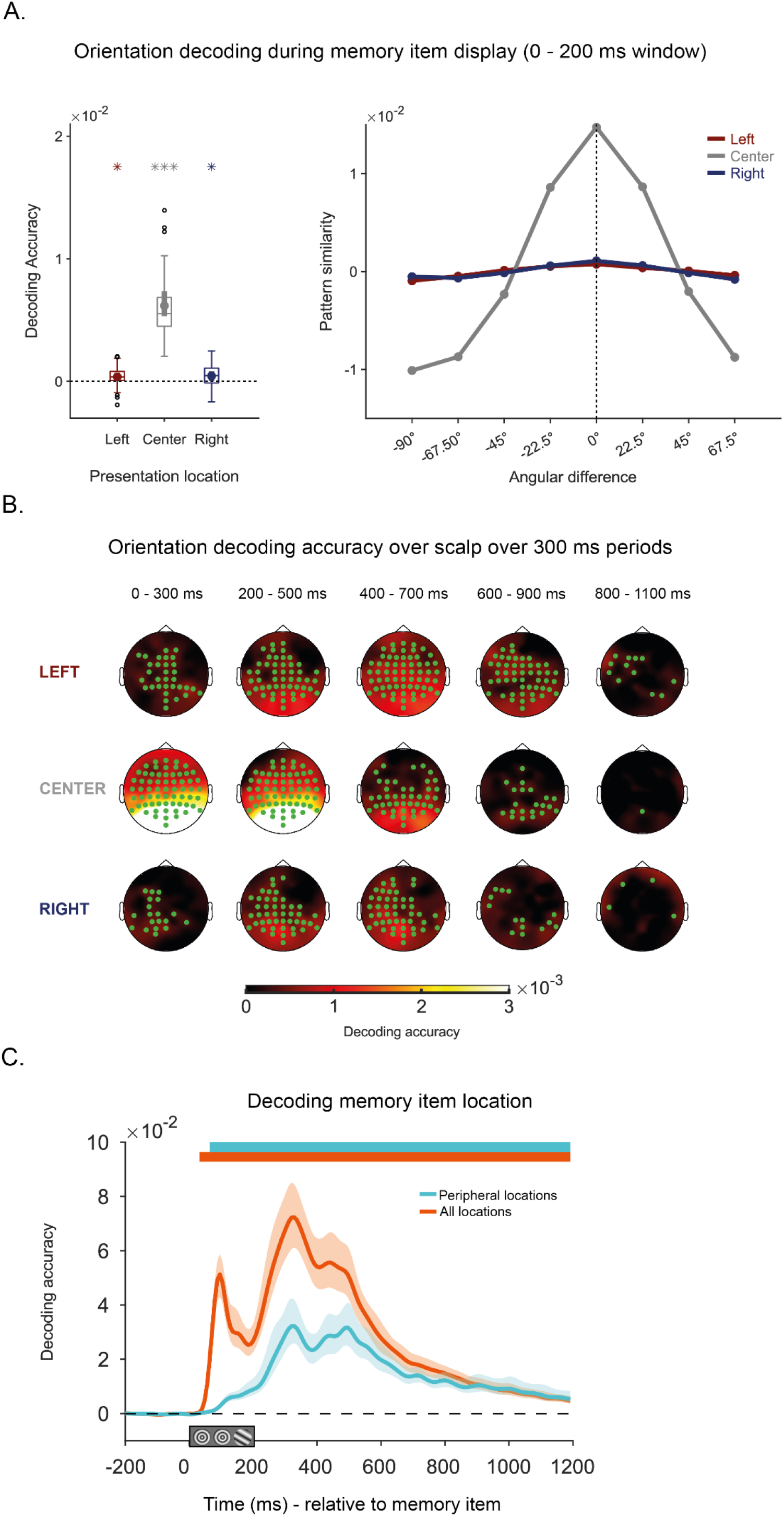
Mean decoding accuracy and similarity curves for item-specific orientations when data is pooled over the entire stimulus encoding period (A). Topographical distribution of electrodes that contribute to accurate decoding of orientation information using searchlight technique (B). Time-course decoding of memory item location (C). *Note.* A. Boxplots showing mean decoding accuracy for items presented on the Left (red), Center (gray) and Right (blue) side of the screen (Left plot). The box borders 25^th^ and 75^th^ percentiles and the whiskers show 1.5 times the interquartile range. The solid line in the middle shows median and the filled dot indicates mean decoding accuracy. Black unfilled circles indicate outliers. Tuning curves reflect pattern similarity as a function of angular deviation from target item, centered to zero (Right plot). B. Electrodes with statistically significant orientation decoding for distinct stimulus locations (green dots, *p* < 0.05, *one-tailed*, n_perm_ = 100 000), when data is pooled over 300 ms periods and decoded in steps of 200 ms. Brighter colours indicate higher decoding accuracy in electrodes at that location. C. Mean decoding accuracy for location of the orientation Gabor, when discriminating across All Locations (orange), and Peripheral Locations (cyan), as a function of time relative to memory item onset. Colored solid lines indicate mean decoding. The shaded zones mark the 95 % C.I.. Horizontal bars at the top mark statistically significant time periods (*p* < 0.05, *two-tailed, cluster corrected*).

In addition, we applied a searchlight technique to determine which electrodes reflected orientation information encoded at distinct locations. The advantage of this technique is that it is searching for the most informative electrodes. The electrodes with above zero decoding (*p* < 0.05, all *one-tailed, n _perm_* = 100 000) are presented in green in Figure 6B, where higher decoding accuracy is reflected by brighter colours. Confirming the time-course analysis, the central item was strongly decodable during sensory encoding in all electrodes, though most strongly in posterior electrodes. Over time, the areas representing the central item shrunk to parietal and occipital electrodes (400 – 700; 600 – 900 ms). Interestingly, peripheral items (Left and Right) were also discriminable during the initial 300 ms, although the effect was quite weak and confined to mostly contralateral electrodes. Here, after stimulus offset, decoding accuracy now actually increased and spread over the scalp, to include frontal and posterior electrodes. Especially during periods with high decoding in the time course analyses (200 – 800 ms, in Fig. 3), peripheral items were represented throughout the scalp (200 - 500 ms, 400 – 700 ms periods), indicating the engagement of brain regions other than just those involved in perception.

Finally, we determined whether we could decode the *location* (rather than the orientation) of the target stimulus from the EEG voltage signal. Note here that the irrelevant locations contained bull’s eye placeholders of the same contrast and spatial frequency, so this analysis allowed us to check if the decoder could at least distinguish the *presence* of a grating if not quite the orientation. When all trials were considered, the location of the task-relevant stimulus (left, right, or center) was decodable shortly after the onset of stimulus (All locations, orange solid line, 34 ms - 1200 ms relative to memory item onset, *p < 0.001*; *two-tailed*, *cluster corrected*). When the analyses were restricted to only the peripheral stimuli, item location could still be distinguished during early processing stages (Peripheral locations, cyan solid line, 64 ms - 1200 ms relative to memory item onset, *p <* 0.001; *two-tailed*, *cluster corrected*). Also noteworthy is that in both cases, location information remained clearly distinguishable throughout the entire epoch, although it was not relevant for the task.

Combined, these findings show that stimulus processing at peripheral locations was not simply absent, or delayed by hundreds of milliseconds. Rather, early signals robustly reflect the presence of an orientation (versus a placeholder), while mustering all the power in our orientation decoding analyses also yields some weak but reliable orientation information. Taken together, we conclude that orientation is registered early at peripheral locations – as would of course be expected – but that it simply cannot be picked up as effectively through EEG. In contrast, signals associated with processes later in the epoch can be picked up in the EEG. We will return to this in the discussion as it is not only methodologically important, but also has important theoretical implications.

## Discussion

Visual working memory is thought to involve visual sensory representations that are maintained through top-down attentional feedback mechanisms. Given that both the sensory input and attentional modulations thereof are known to vary with eccentricity, we investigated the potential consequences of these inhomogeneities for visual working memory, by comparing the electrophysiological correlates for centrally and peripherally presented memoranda.

Behavioral performance indicated that foveally presented orientations were remembered both more frequently (as indicated by reduced guess rates) and with higher precision compared to peripheral items. The higher precision is to be expected given the higher resolution of foveal vision. The increased guess rate for peripheral items is consistent with some reduction in attention for more eccentric stimuli (cf., Van Heusden et al., 2023b; Wolfe et al., 1998), which may result in reduced speed of processing (Staugaard et al., 2016), resulting in additional misses. Note though that while highly reliable, quantitatively speaking the effects were relatively small and memory for the peripheral stimuli was overall remarkably good.

The more surprising then the virtual absence of a decodable orientation signal for peripheral stimuli in the early sensory response of the EEG. While centrally presented orientations became clearly and strongly decodable during the first 200 ms, peripheral orientations were only discernible after mustering additional explorative analyses, and even then only weakly so. Naturally, it is highly implausible that early visual cortex failed to register the peripheral orientations at all, given the fast feedforward connections that characterize it. Rather, it is more likely that the dipoles generated by the early sensory stimulation were undetectable because they were situated deeper into the longitudinal fissure, along the medial walls of the hemispheres (Seki et al., 1996). Importantly, while the failure to detect the early sensory information for peripheral stimuli may be regarded as a shortcoming of our measurements, it actually allows us to draw a number of relevant conclusions, as we will return to below.

In contrast to the early time window, later in the epoch, during the post-stimulus delay period, orientation information was clearly present for peripheral stimuli in the raw voltage signal. This too means that information must have been present during sensory encoding. Furthermore, the dissociation between the early absence and late presence of information confirms that the late component represents a memory-related maintenance process, rather than a remnant of early sensory activity. A number of important conclusions can be drawn from these voltage-based maintenance signals during the delay period: First, the information present was just as strong for the peripheral stimuli as for the central stimuli. Assuming that maintenance involves active feedback mechanisms, there was thus no sign that these mechanisms were weaker for the peripheral memoranda. Second, the orientation information in the raw voltage signal remained location-specific, as there was reliable decoding for each location, but no generalization between locations. This goes against the idea that during this stage, any potential weakness in top-down feedback for peripheral stimuli is simply compensated for by recruitment of other parts of visual cortex, whether through globalization or fovealization. Third, given the observed dissociation between early sensory signals and later maintenance signals – specifically, we observed a localized maintenance signal for central and peripheral stimuli alike, yet failed to observe an early sensory signal for the latter (while observing a very strong sensory encoding signal for the former) – we must conclude that the early sensory input and the later maintenance signals are not making use of the same representations. In other words, the important methodological conclusion here is that successful decoding of orientation from posterior electrodes in the EEG does not as such provide evidence for recruitment of sensory representations. Rather, it appears that these signals must reflect extrastriatal representations. A prime candidate would be a more distributed representation involving parietal cortex, specifically IPS (Bettencourt & Xu, 2016; Christophel et al., 2018; Lorenc et al., 2018), and potentially frontal representations (Lebedev et al.,2004; Passingham & Rowe 2002; Rowe et al 2000; 2005). This is consistent with the emerging distributed activity patterns revealed by our topographic map analysis. We emphasize that the absence of evidence for sensory recruitment of early visual areas in our measurements does not necessarily mean there was no sensory recruitment – just that our observed maintenance signal did not reflect any. However, it is striking that the central patterns then did not generate stronger mnemonic codes, given that these would be able to recruit much more measurable activity in what is presumably a much larger chunk of sensory cortex (Dumoulin & Wandell, 2008; Horton & Hoyt, 1991; Rovamo & Virsu, 1979).

The lack of a generalized voltage signal after stimulus offset runs against the idea of fovealization of peripheral stimuli. Earlier fMRI-based studies have found evidence for foveal representations of peripheral stimuli (Williams et al., 2008) and that this appears to occur within a critical period of around 250 to 500 ms relative to visual encoding (Chambers et al., 2013; Fan et al., 2016). Here, we found no observable generalization within this time period. Several factors might have contributed to the absence of fovealization. First, earlier studies reporting fovealization did not involve foveal stimulation during the presentation of peripheral objects. Conversely, foveal stimulations have been found to disrupt performance for peripheral stimuli whether these central stimulations involved TMS pulses (Chambers et al., 2013) or visual presentations (Fan et al., 2016; Ramezani et al., 2019; Weldon et al., 2016; 2020; Yu & Shim, 2016). Note that in the current design, all locations, including the fovea were stimulated by placeholders (if not targets). While we deemed this necessary to balance initial EEG responses, the activity caused by central placeholders may therefore have prevented or masked foveal representations of peripheral orientations.

While we observed no generalization of orientation information across locations in the voltage signal during earlier part of the maintenance period, we did observe generalization in the later part, including post impulse, and specifically within the alpha band. We believe the transition from location-specific voltage-based code to a location-general alpha-based code reflects a transformation of the memory from a local sensory representation into a functionally relevant representation that can guide subsequent behaviour (Myers et al., 2017; Postle, 2006; van Ede & Nobre, 2023). Specifically, in the current task observers may prepare for the memory test through directing spatial attention or even by pre-activating the direction of the motor response. Thus, “centralized” here does not necessarily mean foveal, but further downstream in the cognitive hierarchy, where sensory recruitment may be replaced by what D’Esposito and Postle (2015) refer to as “sensorimotor recruitment” (see also Olivers & Roelfsema, 2022). Similar changes have been reported earlier in location-bound representations subsequent to the selection and functionalization of memoranda (Kandemir et al., 2022; Panichello & Bushman, 2021), and changes in alpha band activity have been associated with protecting memories against sensory interference, setting attentional priorities within memory, and preparing for retrieval for different tasks (de Vries et al., 2018; Günseli et al., 2019; Pavlov & Kotchoubey, 2022; van Driel et al., 2017). For example, observers may direct covert attention to an internal representation of where they plan to respond. Consistent with this, we found that in the same time period, the orientation became decodable from overt gaze position. Recent work indicates a functional but non-obligatory link between gaze direction and alpha lateralisation (Liu et al., 2022). Note though that while we observed co-occurrence in time between alpha decoding and eye movement decoding strength, we did not observe a direct correlation, suggesting separate origins here. One reason could be that Liu et al (2022) explicitly assessed alpha *lateralisation* in response to stimulus location, while here we measure *content* classification strength.

Finally, an earlier EEG study by Fukuda et al. (2016) also suggested generalization of alpha-based mnemonic signals. Fukuda et al. showed that they could decode a laterally presented orientation not only from alpha power in contralateral electrodes, but also from alpha power in ipsilateral electrodes leading them to claim the occurrence of spatially global representations. While consistent with our general conclusion of there being generalized information in the alpha band, we emphasize that prudence is probably needed in interpreting lateralized activity in this way, because of inherent correlations between ipsilateral and contra-lateral electrodes that do not necessarily reflect a spreading representation. For example, a stimulus-related suppression of alpha power in contralateral electrodes may come with power enhancement in concomitant ipsilateral electrodes (Poch et al., 2017; 2018), thus mirroring the information across the midline. This is not denying the fact that there is substantial other evidence for globalization, as based on the fMRI technique (Ester et al. 2009; Harrison & Tong 2009; Pratte & Tong, 2014; Serences et al. 2009). To our knowledge, our study is the first to provide direct EEG-based evidence tracing the transition from a localized to a generalized memory across peripheral and central positions over time.

To conclude, despite the extensive eccentricity-related inhomogeneities in the visual system itself, we show that visual working memory is remarkably stable and robust across foveal and peripheral locations. We further show that while spatially specific mnemonic representations can be decoded from central and peripheral locations, eventually they merge into a common code that we speculate represents the prospective use, rather than the sensory origin. Finally, we demonstrate that EEG comes with profound limits for decoding early sensory representations of peripheral stimuli, and thus its suitability for testing theories that rely on such representations.

## Data Availability

Analyzed and reported behavioural, EEG and Eye-tracker data, as well as Matlab scripts used to generate Figures can be accessed via this link: (https://osf.io/sv4xt/?view_only=618052fb82bd4dfa91999a9bb956381a)

## Acknowledgements

This project was funded by the Dutch Organization for Scientific Research (NWO; grant 453-16-002, to C.N.L.O.).

## Supplements-1 Decoding orientations at each location from voltage and alpha power

Here we present the time course decoding of orientations specifically for each encoding location, revealing that peripheral item decoding was similar for both eccentric sides (Fig. S1A, Central, gray, 56 ms – 1080 ms relative to memory item onset, p < 0.001; Left, red, 224 ms – 840 ms relative to memory item onset, *p* < 0.001; Right, blue, 240 ms – 408 ms, *p* = 0.037, and 440 ms – 912 ms relative to memory item onset, *p* = 0.005, all *two-tailed*, *cluster corrected*). The matrices (Fig. S1A, right side) reveal a reduction of similarity with increasing distances in angular space for all distinct encoding locations. This parametric-looking pattern provides evidence that the decoding results indeed reflect orientation memory.

Decoding of alpha power was also similar when individual encoding locations were focused on (Fig. S1B, Center, gray, 96 ms to 208 ms, *p* = 0.043, and 624 to 1192 ms relative to item onset, *p* < 0.001; Left, red, 752 ms to 1000 ms relative to item onset, *p* = 0.011; Right, blue, 632 ms to 1192 ms relative to item onset, *p* < 0.001, all *two-tailed*, *cluster corrected*). Similarity matrices again validated that these classifications reflect orientation representations, evident by the gradual decrease in similarity measure with increasing distances to target orientation in angular space.

Voltage data was also decoded following perturbation by the first impulse for orientations at distinct locations. We observed statistically significant decoding only for right peripheral item (Fig. S1C, Right, blue, 216 to 400 ms relative to first impulse onset, *p = 0.019*; Left, red, *p* > 0.05; Center, gray, *p* > 0.05, all *two-tailed*, *cluster corrected*). This effect was not driven by eye movements considering that eye-position decoding only revealed significant results for Left orientation at a different time window (See Fig. S2B). A parametric-looking relationship was reflected in similarity matrix for the Right item, indicating that these representations belonged to orientation memories (S1C, right). Interestingly, for item on the left, similarity matrices suggested a 90 degree shift in neural code, which could explain the negative deflection in mean accuracy in time-course decoding (Fig. S1C, left).

Decoding of orientations after first impulse individually at each encoding location by using alpha band activity yielded weaker but comparable results (Fig. S1D, Center, gray, 112 ms – 1096 ms relative to memory item onset, *p* < 0.001, Left, red, 96 ms – 608 ms relative to first impulse onset, *p = 0.001*; Right, blue, 104 ms – 392 ms, *p* = 0.002, *two-tailed*, *cluster corrected*). Similarity matrices again revealed a parametric-looking relationship between differently oriented memoranda, indicating that impulse driven decoding reflected orientation representations (Fig. S1D, Right).

**Figure S1.**
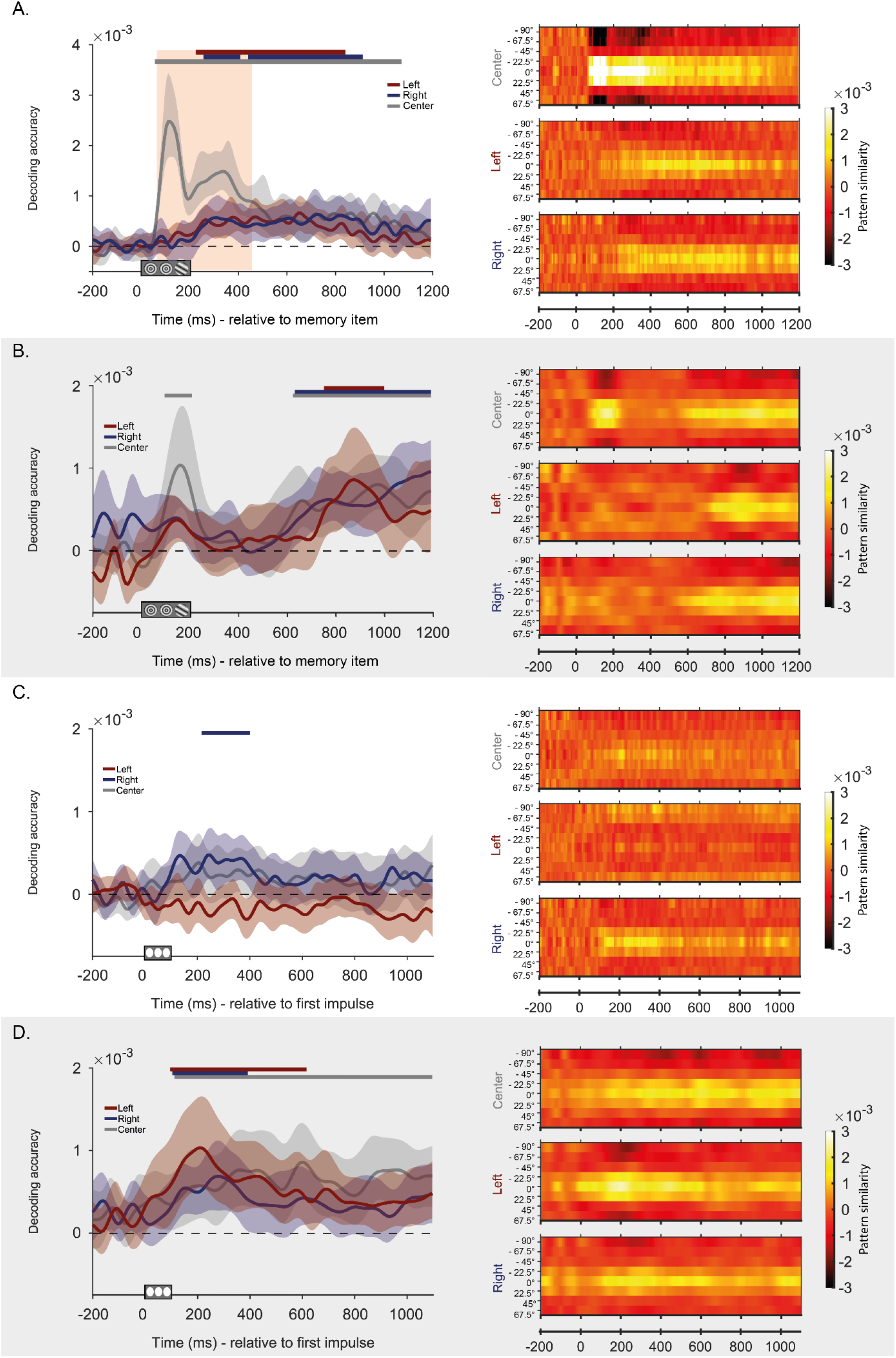
Time course decoding of trial-specific orientations at each location relative to stimulus (A – B) and first impulse onset (C – D) using voltage (A – C) and alpha power (B – D) and respective similarity matrices ()s. recorded by eye tracker, relative to memory item onset (A) and first impulse onset (B). *Note.* A. Mean decoding accuracy for orientations presented at the Center (gray), Left (red) and Right (blue), as a function of time relative to memory item onset, predicted using the voltage values. The purple rectangle marks the time period with statistically significant difference in decoding accuracy between the central item and the average of peripheral items (Left). B. Mean decoding accuracy for orientations presented at the Center (gray), Left (red) and Right (blue), as a function of time relative to memory item onset, predicted using alpha power (Left). C. Mean decoding accuracy for orientations presented at the Center (gray), Left (red) and Right (blue), as a function of time relative to first impulse onset, predicted using the voltage values (Left). D. Mean decoding accuracy for orientations presented at the Center (gray), Left (red) and Right (blue), as a function of time relative to first impulse onset, predicted using alpha power (Left). A-D. Solid lines show mean decoding accuracy averaged over all trials and participants. The shaded zones around the solid lines mark the 95 % C.I.. Horizontal bars at the top mark statistically significant time periods (*p* < 0.05, *two-tailed, cluster corrected*), (Left). A-D. Similarity matrices showing mean centered and reverse-signed Mahalanobis distances between test trials and orientation bins (averaged across all trials and participants), as a function of angular distance to the target item across time (Right).

## Supplements-2 Decoding gaze position for orientations, and correlation with EEG decoding

**Figure S2.**
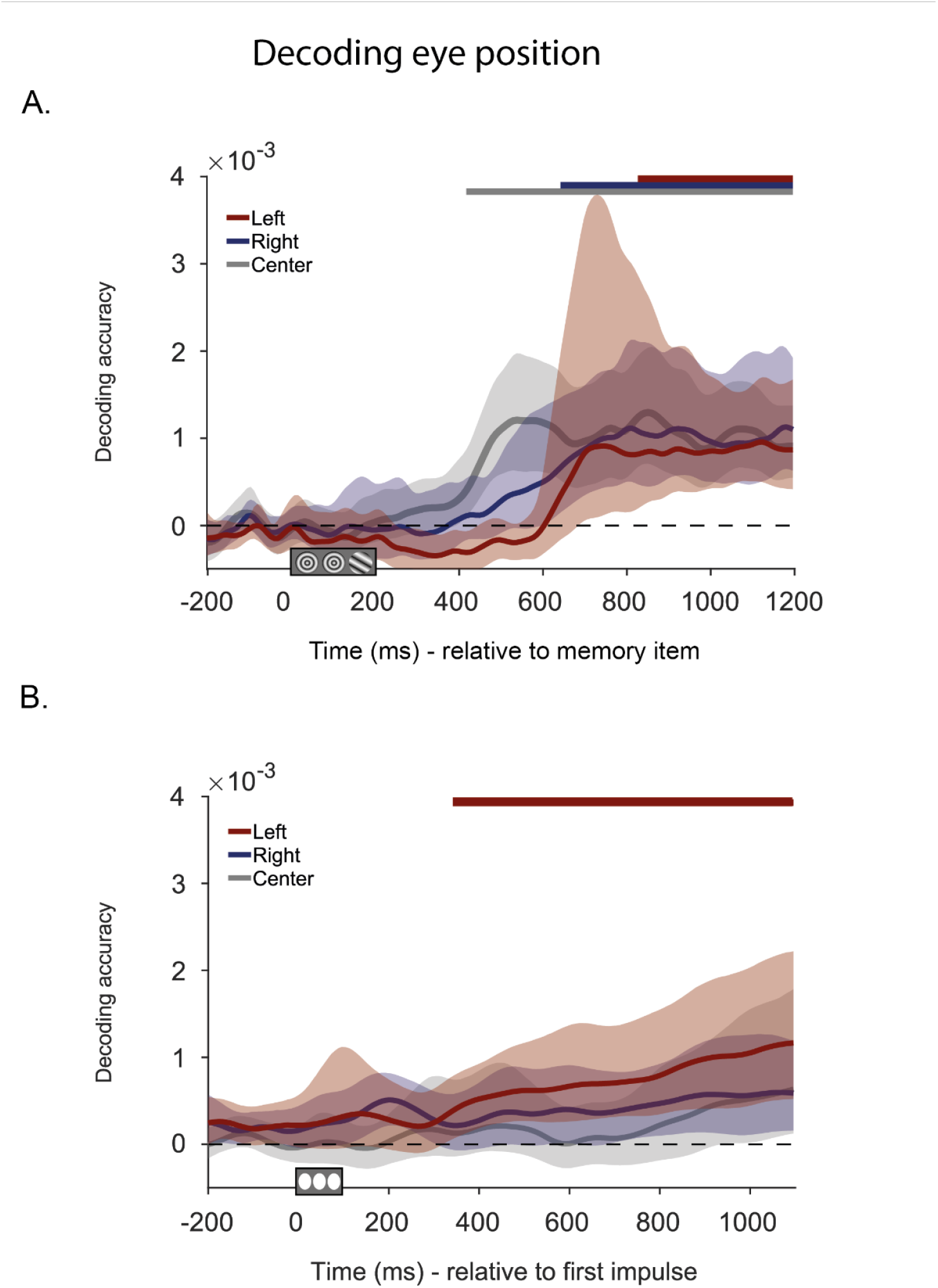
Time course decoding of trial-specific orientations using eye position as recorded by eye tracker, relative to memory item onset (A) and first impulse onset (B). *Note.* A. Mean decoding accuracy for trial specific orientations presented at the Center (gray), Left (red) and Right (blue) trials as a function of time relative to memory item presentation using eye position. B. Mean decoding accuracy for trial specific orientations presented at the Center (gray), Left (red) and Right (blue) trials as a function of time relative to first impulse presentation using eye position. The shaded zones around the lines mark the 95 % C.I. Horizontal bars at the top mark statistically significant time periods (*p* < 0.05, *two-tailed, cluster corrected*).

In order to ensure that signals associated with memories did not originate from gaze position or eye movements, we decoded x-y coordinates of gaze fixations recorded by eye-tracker for trials-specific orientations (Figure S1.A). Statistically significant decoding was observed for all items which emerged following the initial encoding period (Center, gray solid line, 416 ms – 1192 ms relative to memory item onset, *p* < 0.001, Left, red solid line, 824 ms – 1192 ms relative to memory item onset, *p* = 0.007, *two-tailed*, *cluster-corrected*; Right, blue solid line, 640 ms – 1192 ms, *p* = 0.003, *two-tailed*, *cluster-corrected*). These results indicated that gaze position or eye movements correlating with orientation identity emerged later in time, as a side effect of memory maintenance or memory transformation, but initial decodings from voltage measures (e.g., Fig. 3.A.) were not driven by these eye artefacts. Temporal differences in mean decoding between peripheral and central items also support this explanation.

Decoding of gaze position following first impulse yielded significant results only for the peripheral item displayed on the left (Figure S1.B. Left, red solid line, 824 ms – 1192 ms relative to memory item onset, *p* = 0.007, *two-tailed*, *cluster-corrected*). The time-zone with significant decoding did not overlap with significant decoding observed using voltage data, or alpha power. Therefore, we concluded that decoding results following the first impulse were not driven by gaze position or eye movements.

**Figure S3.**
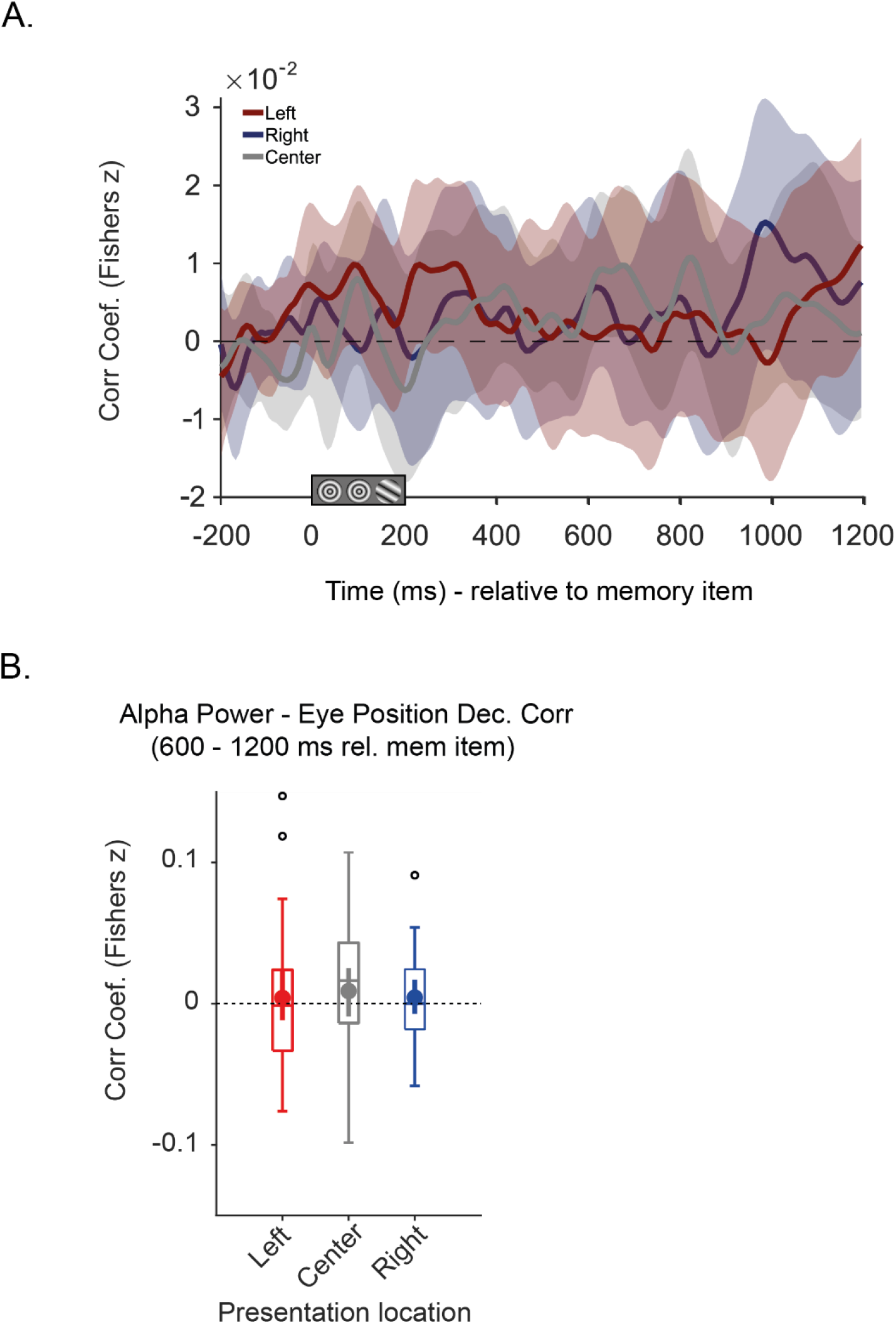
Time course of averaged Fisher’s z transformed correlation coefficients for the correlation of decoding accuracy from gaze position and alpha power at each item location (A), and the boxplot presenting mean Fisher transformed coefficients correlation of alpha power and gaze-position decoding across trials for decoding success within 600-1200 ms period. *Note.* A. Mean Fisher transformed correlation coefficients estimated at each time point using trial-wise decoding accuracy when using alpha power and gaze position for items on Left (red), Center (gray) and Right (blue) side of the screen as a function of time. B. The box borders 25^th^ and 75^th^ percentiles and the whiskers show 1.5 times the interquartile range. The solid line in the middle shows median and the filled dot indicates mean decoding accuracy. Black unfilled circles indicate outliers. B. Tuning curves showing pattern similarity as a function of angular deviation from target item centered to zero. Larger values indicate higher similarity and gradual change indicates parametric-looking relationship between distinct orientations.

Gaze position recorded by eye-tracker could be successfully decoded for trial-specific orientations, and this effect emerged around the same time as alpha power decodings. If these two are related, our evaluation of alpha power decoding would be confounded. The relationship between memory related signal in gaze position and alpha band EEG activity was tested in two steps. First, we assessed the correlation of decoding accuracy from alpha power and gaze position at each time point. The underlying assumption was that if eye movements were the primary determinant of significant classification, then the decoding accuracy obtained from both of these measures would exhibit a positive correlation. The correlation coefficients were estimated at each time point for items at the same location. Fisher transformed coefficients were assessed with cluster-corrected sign-permutation tests, which yielded no significant effects for orientations at any display locations (Figure S2.A). An asynchrony between eye tracker and the amplifier during sampling could have caused this result, by shadowing an actual correlation. Therefore, our next step was assessing the correlation of decoding success over the time window of effect, specifically 600-1200 ms period relative to memory item onset. For this, time course decoding accuracy within 600-1200 ms period was averaged for alpha power and gaze position decoding, providing a single measure in each trial. The correlation coefficient was estimated for each item location for each participant. The Fisher transformed coefficients were tested with group permutation test against zero, which yielded no significant effects (Figure S2.B). We concluded that decoding results for alpha power were not driven by gaze position or eye movements.

## Supplement-3 Temporal Dynamics of Peripheral-Peripheral and Peripheral-Central Generalizations

**Figure S4.**
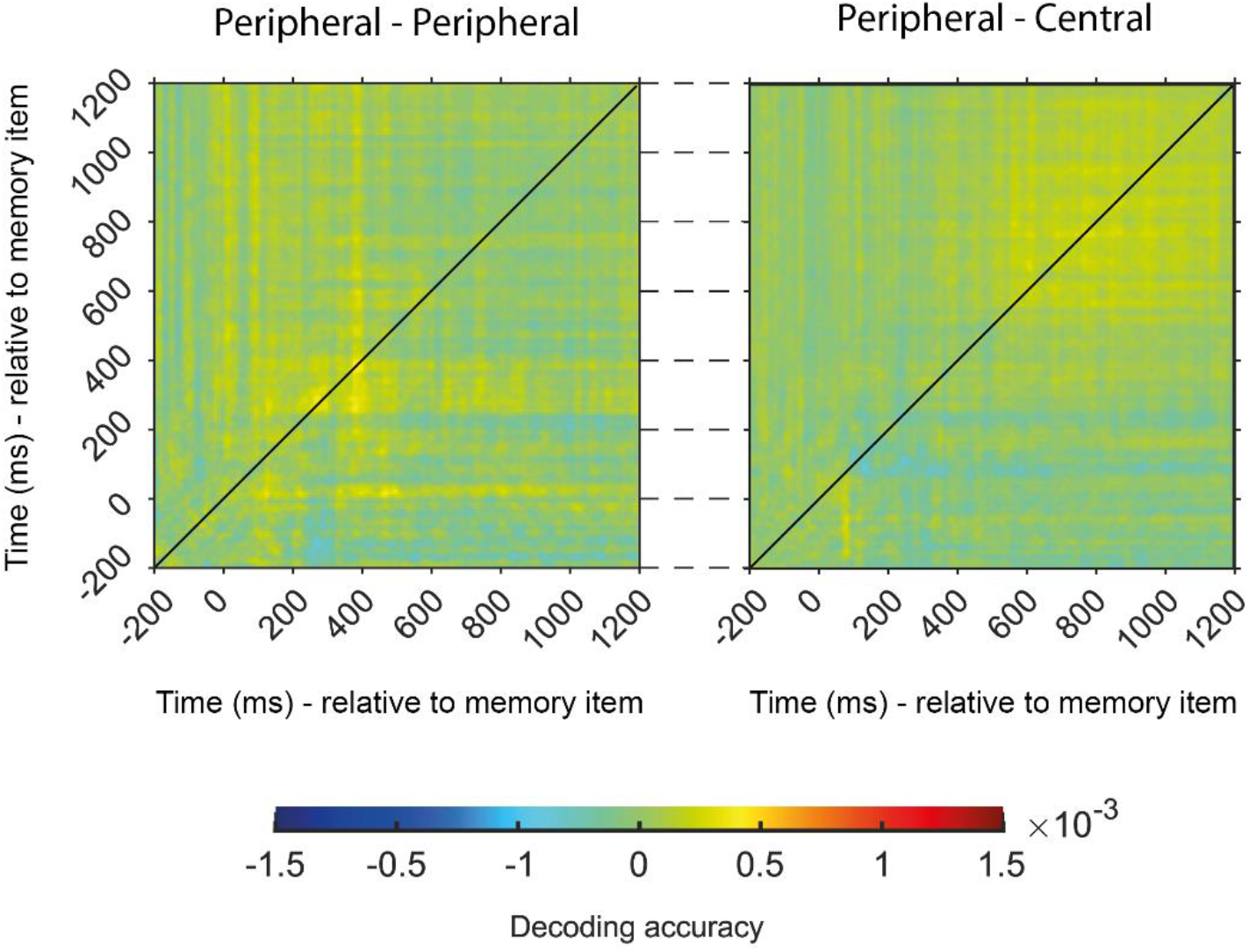
Temporal dynamics of Peripheral-Peripheral (left) and Peripheral-Central (right) generalization of orientation representations across all time points using voltages. *Note.* Temporal dynamics of neural code for Peripheral -Peripheral (Left) and Peripheral-Central (Right) generalization, when data trained at one time point is tested at all time points. Diagonal black line marks the overlap in training and testing time-points. Color intensity reflects decoding accuracy.

The absence of a generalization for neural code across distinct encoding locations could stem from temporal differences in perception. Therefore, we also investigated the temporal dynamics of generalization. For this, we trained our data on orientations at one location at one specific time-point and tested on trials with an item at another location at all time points. A generalization would then be indicated by off-diagonal statistically significant decoding. Across all time-points, we did not observe a Peripheral-Peripheral generalization (Fig. S3, left), or Peripheral-Cental generalization (Fig. S3, right). Importantly, matrices for peripheral-peripheral generalization and peripheral-central generalization revealed different patterns, suggesting that neural code for peripheral items contain some similarities increasing classification success (especially within 200-400 ms period), which is not present for the central item, as Peripheral-Central generalization has increased accuracy in last 600 ms period. This suggests that eccentric items have commonalities that are not shared by centrally encoded items. Overall, these results indicate that memoranda are retained by spatially specific representations.

## Notes

https://osf.io/sv4xt/?view_only=618052fb82bd4dfa91999a9bb956381a

### Competing Interest Statement

The authors have declared no competing interest.

